# Structure, dynamics and roX2-lncRNA binding of tandem double-stranded RNA binding domains dsRBD1,2 of *Drosophila* helicase Maleless

**DOI:** 10.1101/466763

**Authors:** Pravin Kumar Ankush Jagtap, Marisa Müller, Pawel Masiewicz, Sören von Bülow, Nele Merret Hollmann, Bernd Simon, Andreas W. Thomae, Peter B. Becker, Janosch Hennig

## Abstract

Maleless (MLE) is an evolutionary conserved member of the DExH family of helicases in *Drosophila*. Besides its function in RNA editing and presumably siRNA processing, MLE is best known for its role in remodelling non-coding roX RNA in the context of X chromosome dosage compensation in male flies. MLE and its human orthologue, DHX9 contain two tandem double-stranded RNA binding domains (dsRBDs) located at the N-terminal region. The two dsRBDs are essential for localization of MLE at the X-territory and it is presumed that this involves binding roX secondary structures. However, for dsRBD1 roX RNA binding has so far not been described. Here, we determined the solution NMR structure of dsRBD1 and dsRBD2 of MLE in tandem and investigated its role in double-stranded RNA (dsRNA) binding. Our NMR data show that both dsRBDs act as independent structural modules in solution and are canonical, non-sequence-specific dsRBDs featuring non-canonical KKxAK RNA binding motifs. NMR titrations combined with filter binding experiments document the contribution of dsRBD1 to dsRNA binding *in vitro*. Curiously, dsRBD1 mutants in which dsRNA binding *in vitro* is strongly compromised do not affect roX2 RNA binding and MLE localization in cells. These data suggest alternative functions for dsRBD1 *in vivo*.

## INTRODUCTION

In sexually reproducing organisms, the number of X chromosomes differ between males and females. To avoid a lethal imbalance of X-linked gene expression levels, different mechanisms of dosage compensation have evolved (1-3). In *Drosophila* males, the male-specific-lethal (MSL) complex binds with remarkable X-chromosome specificity to PionX sites (4), spreads out along the X-chromosome and achieves two-fold hypertranscription (5). In females, this would be lethal and the RNA binding proteins (RBPs) Sex-lethal (Sxl) and Upstream-of-N-Ras (Unr) repress translation of *msl2* mRNA to prevent formation of the MSL complex (6-8). The MSL complex consists of four core proteins, MSL1, MSL2, MSL3 and males-absent-on-first (MOF) and further accommodates, at least during certain stages of dosage compensation, the helicase Maleless (MLE) and two long non-coding RNAs (lncRNAs), roX1 and roX2 (for ‘RNA-on-the-X’) (2). RoX2 can fold into eight stem loop structures, which we refer to as SL1 to SL8 (**Figure 1A**) (9,10). During assembly of the MSL complex a critical step is the remodeling of roX2 by MLE (10), which is further assisted by Unr (11).

**Figure 1.**
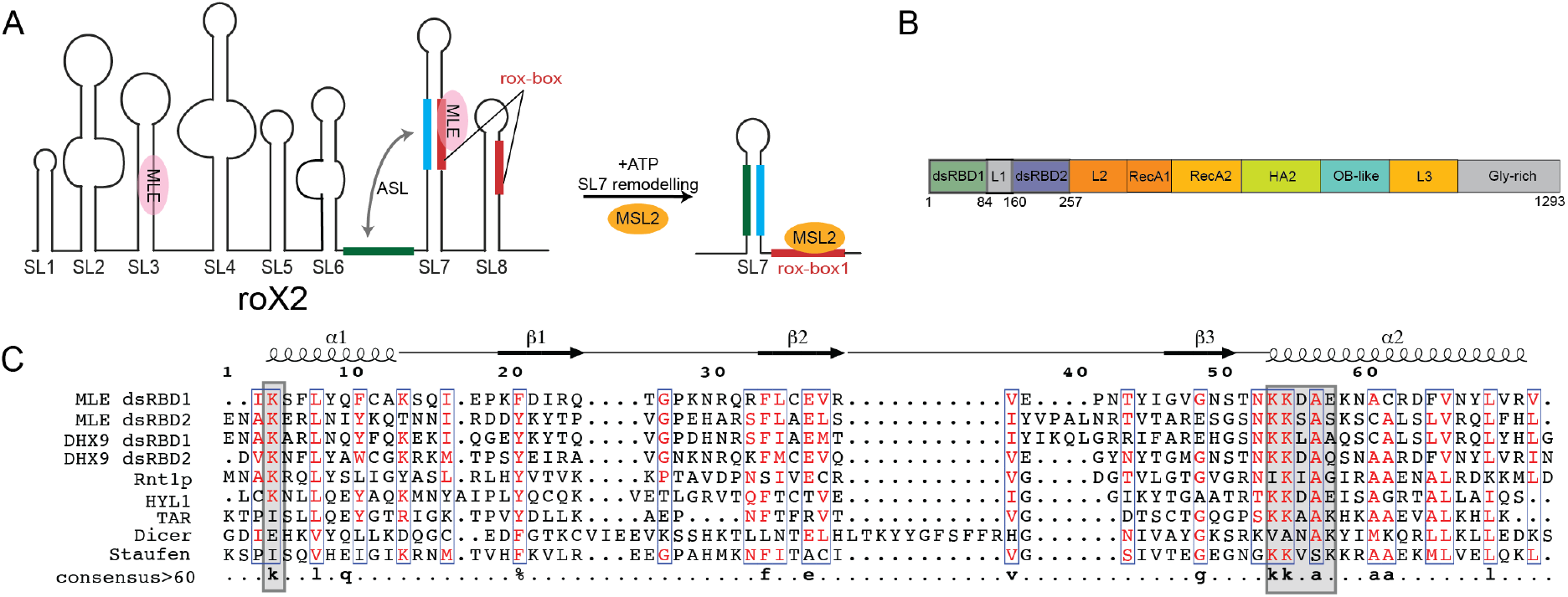
(A) Secondary structure of roX2 RNA consisting of eight stem loops. Stem-loops SL7 and SL8 contain roX-boxes (shown in red). Upon remodelling by MLE, the intervening linker between SL6 and SL7 (green) can base pair with the nucleotides from SL7 (cyan) to form an alternative stem (ASL), thus creating a binding site for MSL2 (9). (B) Domain arrangement of MLE as derived from the MLE crystal structure. (C) Structure based sequence alignment of dsRBD1 and dsRBD2 from MLE with DHX9 dsRBDs (*Homo sapiens*, PDB ID: 3VYX and 3VYY) Rnt1p (*Saccharomyces cerevisiae*, PDB ID: 1T4N), HYL1 (*Arabidopsis thaliana*, PDB ID: 2L2M), TAR RNA binding protein 2 (*Homo sapiens*, PDB ID: 3LLH), DICER (*Schizosaccharomyces pombe*, PDB ID: 2L6M) and Staufen (*Drosophila melanogaster*, PDB ID: 1EKZ). Sequence numbering and secondary structure shown on top with respect to MLE dsRBD1. Consensus sequence with > 60% identity is shown at the bottom of the alignment. RNA binding regions in the dsRBD domains are indicated with grey squares.

MLE is an ATP-dependent RNA helicase of the DExH subfamily of helicases (12). It shares 50% sequence identity with its human homologue DHX9, which has been shown to bind both DNA and RNA (13) and to be involved in diverse cellular functions ranging from DNA replication, microRNA processing, RNA processing and transport, transcription and translation regulation and maintenance of genomic stability (14). Besides its primarily known role in dosage compensation in *Drosophila*, MLE has been shown to be involved in RNA editing and siRNA processing (15,16). During *Drosophila* dosage compensation, MLE has been proposed to bind to SL3 and to a region around SL7 of roX2 to extensively remodel the RNA and to form an alternative stem loop (9,10,17) (**Figure 1A**). MLE’s domain architecture consists of two N-terminal double-stranded RNA binding domains (dsRBDs), followed by the helicase core (RecA1, RecA2, HA2 and OB-fold domains) and a C-terminal glycine-rich region (**Figure 1B**). The structure of the helicase core domains with dsRBD2 has been determined recently (18). Here, dsRBD2 packs against the core domain and is involved in direct roX2 lncRNA binding and essential for localization of MLE to the male X chromosome (18,19). However, there is no structural information regarding dsRBD1 and it has been proposed that dsRBD1 does not bind RNA but is nevertheless involved in X-chromosome targeting (19,20).

In general, dsRBDs are next to RNA recognition motifs (RRM), K-homology (KH) domains and zinc binding domains among the most abundant RNA binding domains (RBDs) (21), which hitherto are known to mainly bind RNA in a structure-specific but not sequence-specific manner involving contacts of the phosphate backbone of an A-form helix. This is mediated usually by helix α1, loop 2 (connecting β1 and β2) and helix α2 that follow the canonical αβββα-fold (21) (**Figure 1C**). This region features a conserved KKxAK motif and binds across the major groove of dsRNA. Sequence specificity in some dsRBDs has been observed, where residues of α1 can contact bases and sugars of the apical loop adjacent to it (22). Also, in ADAR2, a methionine in α1 that protrudes into the minor groove specifies an adenine and replacement by a guanine in the same position abolishes RNA binding (23). However, the question if there is a general sequence-specific recognition code remains unanswered and it is assumed that RNA specificity is based on structure recognition and mediated by other RNA binding proteins as co-factors, which also engage on protein-protein interactions with dsRBDs.

Another specificity determinant has been suggested to be the length of the linker connecting the two dsRBDs. These linkers are often highly flexible and allow the domain to move relative to each other as independent modules, as shown for TRBP, Loqs, and Dicer (e.g. (24-26)). This enables also the probing of different RNA registries shown e.g. for Loqs. The linker between dsRBD1,2 of MLE is 95 residues long, but it is not known whether it is flexible or involved in RNA binding. The length of the linker would allow for reaching across several major grooves of RNA or even between different stem loops.

In the present study, we determined the solution structure of MLE dsRBD1,2 tandem construct and investigated its RNA binding properties and specificity by nuclear magnetic resonance spectroscopy (NMR) and filter binding assays. Furthermore, we investigated the dynamics of the linker in absence and presence of RNA and tested whether the linker has an influence on RNA binding in general. Interestingly, dsRBD1 is clearly involved in RNA binding per se and we could confirm involved residues by mutational analysis. The linker on the other hand is completely dispensable for RNA binding. However, mutations in dsRBD1 that affect dsRNA binding *in vitro* have little effect on binding specific roX RNA in cells and on the localization of the helicase to the dosage-compensated territory, its main place of function, suggesting other functions for dsRBD1 *in vivo*.

## MATERIAL AND METHODS

### Protein cloning, expression and purification

The DNA coding sequence of dsRBD1 (M1-G84), dsRBD2 (F155-D257) and tandem dsRBD1,2 (M1-D257) were amplified from the full *mle* gene and cloned into the N-terminal His6-tag and TEV protease cleavage site containing expression vector pETM11 using the restriction-free approach. The mutants: K4E, K53E, K54E, K53+K54E, K225E and K54E+K225E in the dsRBD1, dsRBD2, dsRBD1,2 or full length MLE context were generated by site-directed mutagenesis using a QuickChange XL kit (Ambion). For the linker deletions, dsRBD2 plus the desired linker constructs were amplified and cloned into the pETM11-dsRBD1 construct using the restriction-free approach.

DsRBD constructs were expressed in *E. coli* BL21 (DE3) cells by addition of 0.3mM IPTG at OD_600_=0.9 and overnight growth at 18°C. Next, the cells were collected, resuspended in lysis buffer (50mM NaPi, 300mM NaCl, 20mM imidazole, 2mM β-mercaptoethanol, pH 7.5), lysed by sonication and spun down. The supernatant was loaded onto a HisTrap HP column (GE Healthcare) and eluted with an imidazole gradient in buffer containing 50mM NaPi, 300mM NaCl, 300 mM imidazole and 2mM βmercaptoethanol, pH 7.5. His6-tag was removed by addition of TEV protease and simultaneous dialysis into low salt buffer (20mM NaPi, 200mM NaCl, 1mM DTT, pH 7.5) overnight at 4°C. Next, the sample was applied to a second affinity chromatography using a HisTrap HP column and the flow-through fraction was further loaded onto a HiTrap Q and HiTrap Heparin column (GE Healthcare) in order to remove bound bacterial RNA. The protein was eluted with a salt gradient in the elution buffer (20mM NaPi, 1M NaCl, 1mM DTT, pH 7.5) and applied to size-exclusion chromatography on the HiLoad 26/600 Superdex S75 pg column (GE) equilibrated with NMR buffer (20mM NaPi, 200mM NaCl, 1mM DTT, pH 6.5).

For NMR experiments, the proteins were produced with different labelling schemes. For ^13^C and/or ^15^N labelling, the proteins were expressed in M9 minimal media containing 2g/l 13C-labelled glucose and/or 0.5 g/l ^15^N-labelled ammonium chloride. Uniformly deuterated proteins were obtained by expression in M9 media containing 99.8 % D_2_O (Sigma). To final NMR samples, 10 % D_2_O and 0.02 % NaN_3_ were added for the deuterium lock and to prevent bacterial growth.

Wild type and mutant full-length MLE was expressed in SF21 insect cells using recombinant baculoviruses and purified by FLAG-affinity chromatography as described (18).

### RNA production

SL7^14merLoop^ RNA (5’-GUG UAA AAU GUU GCA AUA UAU AGU AAC GUU UUA CGC-3’) was prepared by *in vitro* transcription using T7 polymerase produced in-house, unlabelled rNTPs (Sigma-Aldrich), and a synthetic DNA template. SL7^long^ RNA (5’- ACUGAAGUCUUAAAAGACGUGUAAAAUGUUGCAAAUUAAGCAAAUAUAUAUGCAUAUAUGGGUAACGUUUUACGCGCCUUAACCAGU-3’) was produced the same way, except the template was cloned into pUC19 plasmid DNA and contained a hammerhead ribozyme (HH) cleavage site (in *cis*) at the 5’ end and a VS (Varkud satellite) ribozyme recognition sequence at the 3’ end for cleavage in *trans*. After transcription, proteins were removed by phenol/chloroform extraction. The RNA was purified by denaturing 12% PAGE and extracted from the gel by electro-elution. The final sample was concentrated and dialyzed against 20mM NaPi, pH 6.5 buffer.

The SL7^18mer^ and SL7^23mer^ RNA duplexes were obtained by annealing of two purchased RNA oligos (IBA, Göttingen): SL7^18mer^_up (5’-AGACGUGUAAAAUGUUGC -3’) with SL7^1^8mer_bot: (5’-GUAACG UUUUACGCGCCU-3’) and SL7^23mer^_up (5’-UUAAAAGACGUGUAAAAUGUUGC-3’) with SL7^23mer^_bot (5’-GUAACGUUUUACGCGCCUUUUCC-3’) respectively. Another UR^23mer^ dsRNA, used in a control experiment, was prepared the same way using UR^23mer^- up (5’-UAAAGUGCUUAUAGUGCAGGUAG-3’) and UR^23mer^- bot (5’-ACCUGCACUAUAAGCACUUUAAG-3’).

### NMR data acquisition and structure determination

All spectra were acquired using Avance III Bruker NMR spectrometers with proton Larmor frequencies of 600 MHz, 700 MHz or 800 MHz at 298 K, equipped with cryogenic (600 and 800 MHz) or room temperature (700 MHz) triple resonance gradient probe heads. For structure determination data were recorded with 0.8 mM protein in 20 mM sodium phosphate (pH 6.5), 200 mM NaCl and 1 mM DTT with 10% D2O added for the deuterium lock. All spectra were processed using NMRPipe (27) and analysed using CCPNmr (28). Protein backbone assignments were obtained from HNCO, HNCA, CBCA(CO)NH, HNCACB (29) and HBHA(CO)NH (30) experiments. HCCH-TOCSY and ^15^N-/^13^C-edited NOESYHSQC experiments were used for amino acid side chain resonance assignments. 2D ^1^H–^13^C HSQC, HBCBCGCDHD, HBCBCGCDCEHE (31) and ^15^N-/^13^C-edited NOESY-HSQC spectra were used for side chain resonance assignments of aromatic residues. Distance restraints were obtained from ^15^N-/^13^C-edited NOESY-HSQC spectra.

The NMR ensemble was calculated using CYANA 3.97 (32) based on automated NOE crosspeak assignment and torsion angle dynamics. Automatically assigned NOEs and completeness of NOE crosspeak assignments were manually inspected for validation. Distance restraints from the CYANA calculation and TALOS+-derived dihedral angle restraints (33) were used in a water refinement calculation using ARIA 1.2 (34). Structures were validated with PROCHECK (35). Structural statistics are shown in **Table 1**. Molecular images were generated with PyMol (Schrödinger).

**Table 1.** Structural statistics for MLE-dsRBD1,2.

| Structural statistics for MLE-dsRBD1,2<sup>a</sup> |  |
| --- | --- |
| Distance restraints |  |
| Total NOEs | 2734 |
| Intra-residue | 761 |
| Sequential ($\vert i - j\vert = 1$) | 870 |
| Short-range, $\vert i - j\vert \leq 1$ | 1631 |
| Medium-range ($\vert i - j\vert \leq 5$) | 385 |
| Long-range ($\vert i - j\vert > 5$) | 718 |
| Hydrogen bonds | 103 |
| Dihedral restraints ($\phi + \psi$) | 782 |
| Quality analysis (mean ± standard deviation) | |
| Distance restraints (Å) | 0.032 ($\pm 0.006$) |
| Dihedral angle restraints (°) | 0.89 ($\pm 0.32$) |
| Deviation from idealized geometry |  |
| Bond length (Å) | 0.003 ($\pm 0.0004$) |
| Bond angles (°) | 0.52 ($\pm 0.017$) |
| Impropers (°) | 1.71 ($\pm 0.13$) |
| Average pairwise r.m.s. deviation (Å)<sup>a</sup> |  |
| Backbone (dsRBD1/dsRBD2) | 0.59/0.87 |
| Heavy (dsRBD1/dsRBD2) | 1.17/1.46 |
| Ramachandran values (%)<sup>a,b</sup> |  |
| Most favoured regions | 92.2% |
| Allowed regions | 7.8 % |
| Generously allowed regions | 0 % |
| Disallowed regions | 0 % |

<sup>a</sup> For residues 5-26, 35-73 (dsRBD1) and 170-193, 202-243 (dsRBD2)
<sup>b</sup> Using PROCHECK

### NMR titration

NMR titrations for dsRBD1,2 were performed at 0.3 mM protein concentration. ^15^N labelled dsRBD1,2 was titrated with various ratios of different RNA oligos described in the results section and monitored by recording ^1^H,^15^N HSQC spectra. For SL7^14merLoop^ titrations, deuterated ^15^N labelled dsRBD1,2 protein was used. Titrations for the single domains, 0.05 mM dsRBD1 wt or mutants or 0.1 mM dsRBD2 were titrated with 18-mer double stranded roX-box RNA derived from SL7 respectively. Data analysis for calculating chemical shift perturbations and dissociation constants were performed with the CCPN analysis software (28).

### NMR relaxation

Measurements of *R_1_, R_2_* and {^1^H}-^15^N heteronuclear NOE experiments for dsRBD1,2 and {^1^H}-^15^N heteronuclear NOE experiments for dsRBD1,2 + 1.2x SL7^14merLoop^ RNA were acquired at proton Larmor frequencies of 600 MHz and 800 MHz, respectively, at 298 K using standard pulse sequences (36,37). For the *R_1_* experiment for dsRBD1,2, relaxation delays of 20, 50, 100, 150, 250, 400, 500, 650, 800, 1000, 1300, and 1600 ms were used. Duplicate data points for estimation of uncertainties in peak volumes were recorded for the 20 ms relaxation delay. For the *R_2_* experiment, relaxation delays of 16, 32, 48, 64, 80, 96, 112, 128, 160, 192, 224, and 256 ms were recorded. Duplicate data points were recorded for the 16 ms relaxation delay (36). PINT (38,39) was used for peak integration and data fitting to derive spin relaxation parameters.

### RNA filter binding assays

The reaction buffer for the binding assay was 20 mM HEPES/NaOH pH 8.0, 150 mM NaCl, 5 mM MgCl_2_. The reaction volume was 200 μl. Structured RNA probes were refolded prior to experiments by heating at 85°C for 3 minutes and slow cooling to room temperature. Increasing concentrations of the protein were incubated with 5’-labelled RNA or oligoribonucleotides for 10 minutes on ice. The mixture was then filtered through Whatman 0.45 μm pore size nitrocellulose filters. Only the RNA complexed with protein was retained on the filters and detected by scintillation counting. Since the amount of RNA used was extremely small (0.03 pmol of molecules, 1.5-2×10^4^ cpm), the protein concentration at 50% retention corresponds to the apparent dissociation constant *K_D_*. Data fit and K_D_ calculations were performed with GraphPad Prism 8 using a single site saturation binding model.

### Cell culture and RNA immunoprecipitation

*Drosophila* S2 cells were cultured in standard conditions. Stable S2 cell lines expressing wild type or mutant MLEfl fused to C-terminal GFP were generated as described (18). Point mutations in pHsp70-MLEfl-eGFP plasmids were introduced using site-directed mutagenesis using a QuickChange XL kit (Ambion). Briefly, for generation of stable cell lines, 2 µg pHsp70-ML_FL_-eGFP wild type or mutant plasmid was co-transfected with 0.1 µg plasmid encoding a hygromycin resistance gene using the Effectene transfection reagent (Qiagen). Stable MLEfl-GFP expressing clones were selected in complete medium containing 0.3 mg/ml hygromycin for a duration of four weeks.

Native RNA immunopreciptation (RIP) of MLE-GFP and mutant derivatives was performed as described in (18) with slight modifications. For each replicate, 1 x 10^8^ exponentially grown S2 cells expressing MLE-GFP were used. Non-transfected S2 cells served as negative control (mock). Cells were lysed in 1 ml of lysis buffer (20 mM HEPES pH 7.6, 125 mM NaCl, 0.05 % SDS, 0.25 % sodium deoxycholate, 0.5 % NP40, 2 mM MgCl_2_) supplemented with 0.05 U/µl RNase-free recombinant DNase I (Roche), 0.4 U/µl RNasin recombinant RNase inhibitor (Promega) and 1x complete EDTA-free protease inhibitor (Roche). Of the lysate, 5% was kept as input material for RNA quantification and protein analysis. The lysate was mixed with 30 µl GFP-Trap beads (Chromotek) and incubated at 4°C for 2 hours on a rotating wheel. Beads were washed three times with RIP-100, RIP-250, RIP-100 buffer (25 mM HEPES pH 7.6, 0.05% NP40, 3 mM MgCl_2_ with 100 mM NaCl and 250 mM NaCl, respectively). RNA was extracted as described (18) from 75% of bead material. GFP-pull down efficiency was controlled using anti-MLE western blot analysis of 25% bead material. RNA samples (Input and IP) were analyzed by quantitative reverse-transcription PCR (qRT-PCR) with SYBR green dye (Applied Biosystems) using primers specific for *roX2* and *7SK* (**Table S1**). RNA enrichment of MLE-GFP and its mutants was calculated as IP/Input and represented relative to WT MLE. Highly expressed *7SK* RNA served as an MLE-unbound control in each experiment.

### Immunocytochemistry

Immunolocalization assays were performed essentially as described before (18). Briefly, stable cell lines of *Drosophila* S2 cells expressing MLE-GFP and its mutants were used. Cells at a density of 3 x 10^6^/ml were immobilized on coverslips by settling for 20 min and then fixed with PBS/3.7% formaldehyde. The cells were then permeabilized with PBS/0.25% Triton X-100 for 6 mins on ice and blocked with Image iT^FM^ FX signal enhancer for 30 min (40). The fixed and permeabilized cells were then incubated with primary antibodies against GFP (mouse; Roche), MLE (rat) (19) and MSL2 (rabbit) (41) at 4°C overnight. After washing with PBS/0.1% Triton X-100, fluorophore coupled secondary antibodies against mouse (Alexa Fluor 488), rat (Cy3) and rabbit (Alexa Fluor 647) were added for 1 h at room temperature. DNA was stained with DAPI. After washing with PBS/0.1% Triton X-100, cells were mounted in VECTASHIELD.

Images were acquired at the Core Facility Bioimaging of the Biomedical Center of the LMU using a Leica DMi8 widefield fluorescence microscope with 63x oil immersion objective NA = 1.4. The pixel size was 102 nm. The effective excitation and emission filter combinations were as follows (Ex/Em): DAPI: 379-401 nm / 450-490 nm, GFP/Alexa Fluor 488: 458-482 nm / 500-550 nm, Cy3: 543-551 nm / 565-605 nm, Alexa Fluor 647: 625-655 nm / 670-770 nm. Nuclei and X-territory segmentation was performed in CellProfiler (42) using DAPI and anti-MSL2 staining respectively. MLE-GFP non-expressing cells and cells exhibiting strong MLE-GFP overexpression were not included in the analysis. The upper and lower thresholds were extracted from whole cell population GFP intensity plots that highlighted non-expressing, moderate and highly expressing cells. This analysis was performed with two independent biological replicates. The enrichment of MLE-GFP on MSL2-stained X-territories was measured by calculating the ratio of the intensity of MLE-GFP in the MSL2 territory vs in the whole nucleus.

## RESULTS

### The tandem dsRBD domains of MLE are independent structural modules

First, we wanted to assess whether the two dsRBD domains interact with each other despite the long 95-residue linker. Interdomain interactions have been reported for other RNA binding domains even in absence of RNA and could constitute a determinant of specificity by conformational selection (43,44). Therefore, we recorded ^1^H,^15^N-HSQC spectra of the individual dsRBD domains and compared them with the spectrum of the tandem dsRBD1,2 construct (**Figure 2A**). The three spectra are very well superimposable and the chemical shifts of the individual dsRBD domains are very similar to the tandem dsRBD1,2 except for the expected changes for residues at respective termini caused by the changed chemical environment of the additional peptide bonds (**Figure 2B**). Thus, both domains do not seem to interact with each other in solution. For further evidence, we determined the rotational correlation times (τ_c_) of the individual domains within the tandem construct by measuring the ^15^N NMR relaxation parameters *R_1_* (longitudinal relaxation rate) and *R_2_* (transverse relaxation rate) (**Figure S1**) and from these determined the residue-wise and the average τ_c_ of the two domains (**Figure 2C**) assuming isotropic tumbling in solution for each domain (45). The average τ_c_ of dsRBD1 and dsRBD2 is 9.2 ns and 11.7 ns, respectively. These are expected values for domains of this size when tumbling independently but being attached to a long flexible tail or linker. For a 29 kDa protein and assuming both domains would interact in solution to form a single module, a much higher τ_c_ of ~17 ns would be expected. However, as the two domains show different and considerably lower τ_c_ than 17 ns, we conclude that the two domains do not tumble together and act as independent structural modules in solution.

**Figure 2.**
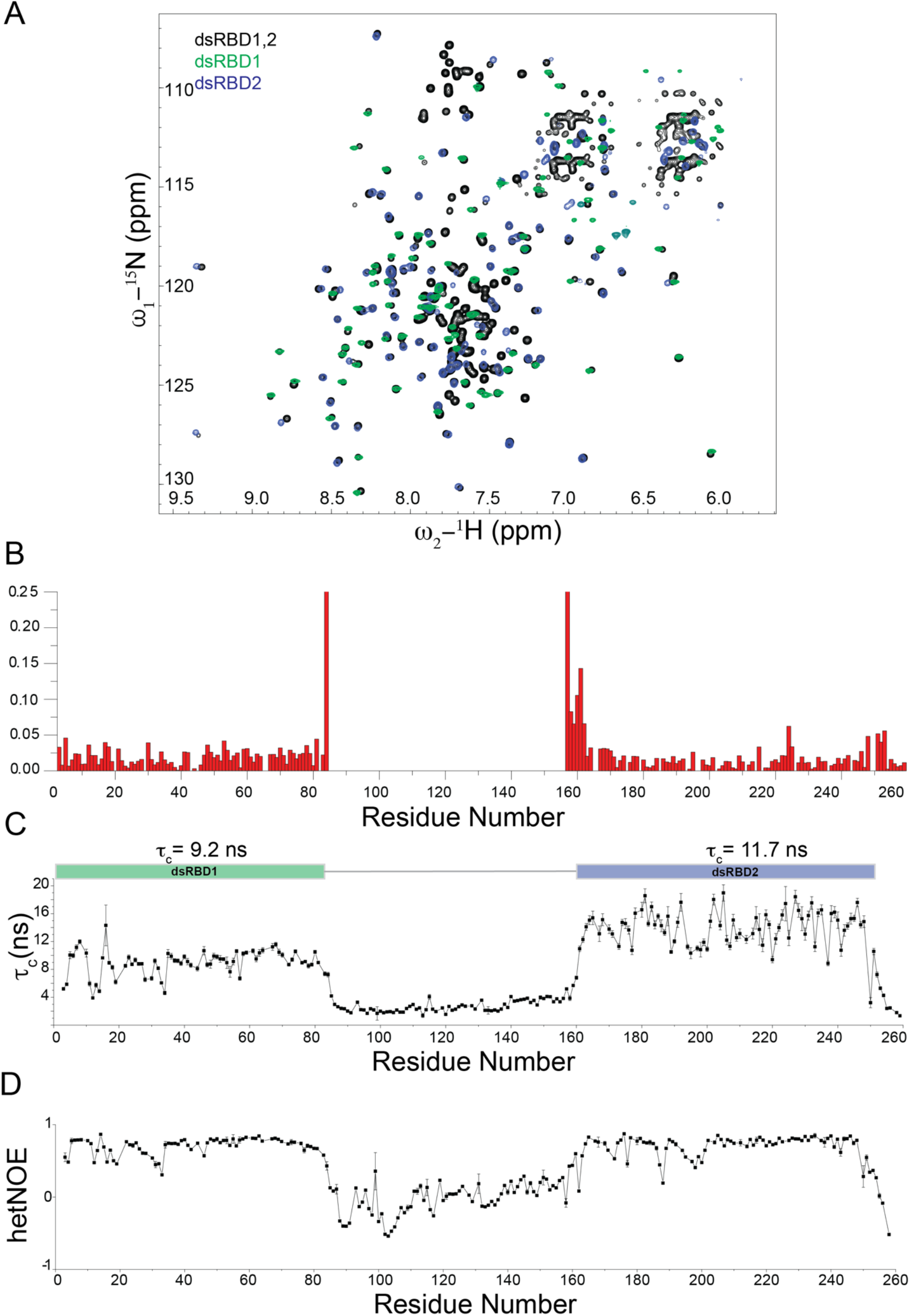
(A) ^1^H,^15^N-HSQC NMR spectrum of dsRBD1,2 (black), dsRBD1 (green) and dsRBD2 (blue) are shown. No major chemical shift differences between the three spectra could be observed suggesting that the two domains act as independent modules in solution. (B) Chemical shift perturbations in dsRBD1,2 compared to individual domains. Residues located at the C-terminus of dsRBD1 and the N-terminus of dsRBD2 are the only ones, which exhibit chemical shift differences, but this derives from being the terminal residues in case of individual domains and being connected to the linker in context of the dsRBD1,2 construct. (C) Rotational correlation times (τ_c_) for individual residues as calculated from the ratio of ^15^N *R_2_/R_1_* relaxation rates are shown. Average τ_c_ for each individual domain is shown on top of the graph. Error bars are calculated from duplicate relaxation delays (see methods). (D) {^1^H}–^15^N heteronuclear NOE values for the MLE-dsRBD1,2. Both graphs visualize the different dynamics of the linker region compared to the domain regions. The linker region is highly dynamic.

In addition to this, dsRBD2 exhibits a higher τ_c_ than dsRBD1. This would be expected, if dsRBD2 has a slightly higher molecular weight due to its extension of ordered residues at either N or C termini or due to an aggregation tendency with itself, which would cause an apparent increase in its molecular weight. In order to determine if dsRBD2 has more ordered residues then dsRBD1, we recorded {^1^H}–^15^N heteronuclear NOE experiments (**Figure 2D**). These experiments clearly show that residues 5-80 (76 residues) and 164-248 (85 residues) are well ordered and correspond to a calculated molecular weight of 8.8 kDa and 9.7 kDa for dsRBD1 and dsRBD2, respectively. Therefore, the increased τ_c_ for dsRBD2 can be explained by the larger size of the dsRBD2 domain.

### Solution structure of the dsRBD1,2 tandem domain construct

To understand the structures of both dsRBDs in their native arrangement, we determined the solution structure of tandem dsRBD1,2. To this end, we obtained NMR resonance assignments (backbone and side chain) to a completion of 90.5%, which enabled automated NOE assignment and structure calculation. The structures of the individual dsRBDs within the dsRBD1,2 tandem construct are shown in **Figure 3** along with the superposition of the 10 lowest energy structures after water refinement. Structural statistics are listed in **Table 1**. The NMR ensemble of each domain converged with a backbone RMSD of 0.59 Å and 0.87 Å and heavy atom RMSD of 1.17 Å and 1.46 Å for dsRBD1 and dsRBD2, respectively (**Figure 3A, B, Table 1**). Both domains show the typical dsRBD fold with residues 4-74 and 170-245 for dsRBD1 and dsRBD2, respectively forming an α-β-β-β-α fold consisting of three anti-parallel b-sheets covered on one side by two a-helices (**Figure 3C, D**).

**Figure 3.**
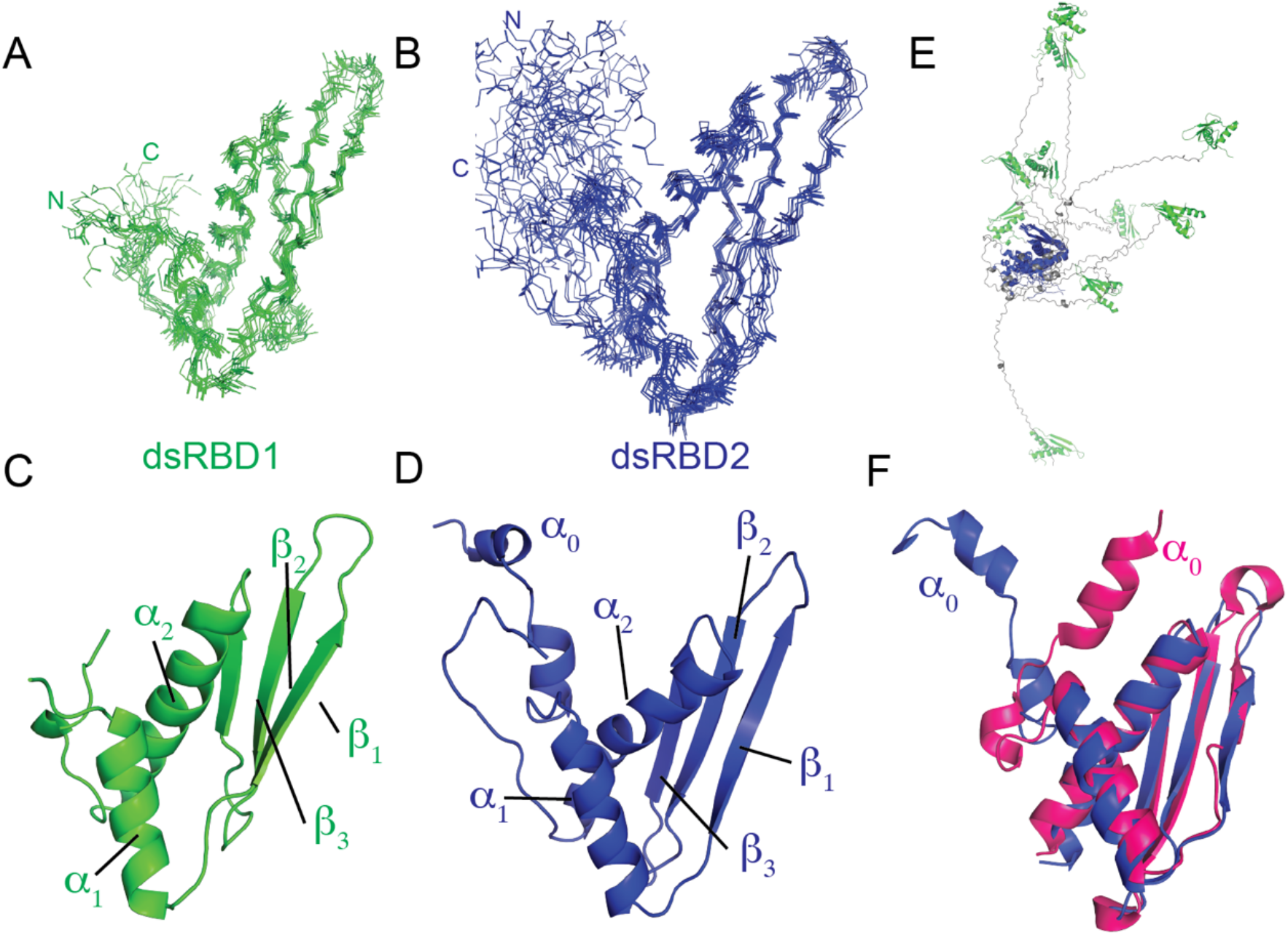
NMR ensemble of 10 lowest energy structures generated by backbone superposition of (A) dsRBD1 residue 1-84 and (B) dsRBD2 (residue 147-259) in the MLE-dsRBD1,2 NMR structure. The N- and C- termini of the two domains are marked for clarity. Cartoon representations of (C) dsRBD1 and (D) dsRBD2 are shown along with the labelled secondary structure elements. (E) Superposition of dsRBD2 (blue) backbone in the dsRBD1,2 ensemble shows that dsRBD1 (green) does not adopt any fixed orientation relative to dsRBD1 owing to the lack of NOE’s between the two domains. Linker residues are shown in grey. (F) Comparison of the solution structure of dsRBD2 presented in this work (blue) and the crystal structure of dsRBD2 as part of MLE_core_ (PDB ID 5AOR) (magenta). The major difference is the orientation of helix α_0_ with respect to the dsRBD domain. However, our NMR relaxation data and absence of NOEs between α_0_ and domain residues indicate that in the absence of other MLE domains this helix is dynamic.

The two dsRBD domains also superimpose well with an RMSD of 1.1 Å. Structural superposition of the two domains showed that the α_1_ helix is displaced by one helical turn; a feature observed previously in other tandem dsRBD domains (46) and the linker connecting β_2_-β_3_ sheets of dsRBD2 is longer by four residues compared to the dsRBD1 domain (**Figure S2A**).

The linker region comprising residues 75 to 169 does not give rise to any long-range NOEs and remains disordered in structure calculations. The highly flexible nature of the linker has been already confirmed with NMR relaxation experiments in the previous section. Thus, both domains do not have a fixed orientation with respect to each other in solution (**Figure 3E**). The flexible linker allows both domains to theoretically span a distance of ~150 Å, which corresponds to reaching across ~4 major grooves of A-form dsRNA.

We can further confirm the formation of an additional helix α0 in solution, which is also present in the crystal structure of the MLE helicase core domain and dsRBD2 (18) (PDB ID: 5AOR) (**Figure 3F**). However, this helix is highly dynamic as can be seen from relaxation analysis and the lack of NOEs between this helix and other parts of the domain confirm this flexibility.

### RNA binding of individual dsRBD domains

Sequence alignment of dsRBD1 and 2 revealed that both domains contain conserved RNA binding residues typically involved in dsRBD-RNA interactions (21) (**Figure 1C**). Therefore, we found it surprising that only dsRBD2 had been identified to bind RNA (19). To determine the contribution of each dsRBD domain for RNA binding, we performed ^1^H,^15^N-HSQC NMR titrations with each individual dsRBD domain and dsRNA (**Figure 4, S3**). Uridine-rich stem-loop structures and single-stranded polyU sequences have been shown as preferred substrates for MLE (9-11) and remodeling of the uridine-rich roX-box regions in roX RNA has been shown to recruit MSL2 to the MLE-roX2 complex, suggesting a role of MLE during initial assembly of the MSL complex (10,17) (**Figure 1A**). To study the interaction of MLE dsRBDs with RNA, we chose RNAs derived from roX-box of roX2 stem loop 7 (SL7). A minimal dsRNA length of ~10-12 base pairs (bp) has previously been reported sufficient for the binding of individual dsRBDs to dsRNA (23,46,47). Therefore, for RNA titration of individual dsRBD domains, an 18 bp long roX-box RNA (SL7^18mer^) was used (**Figure S4**).

**Figure 4.**
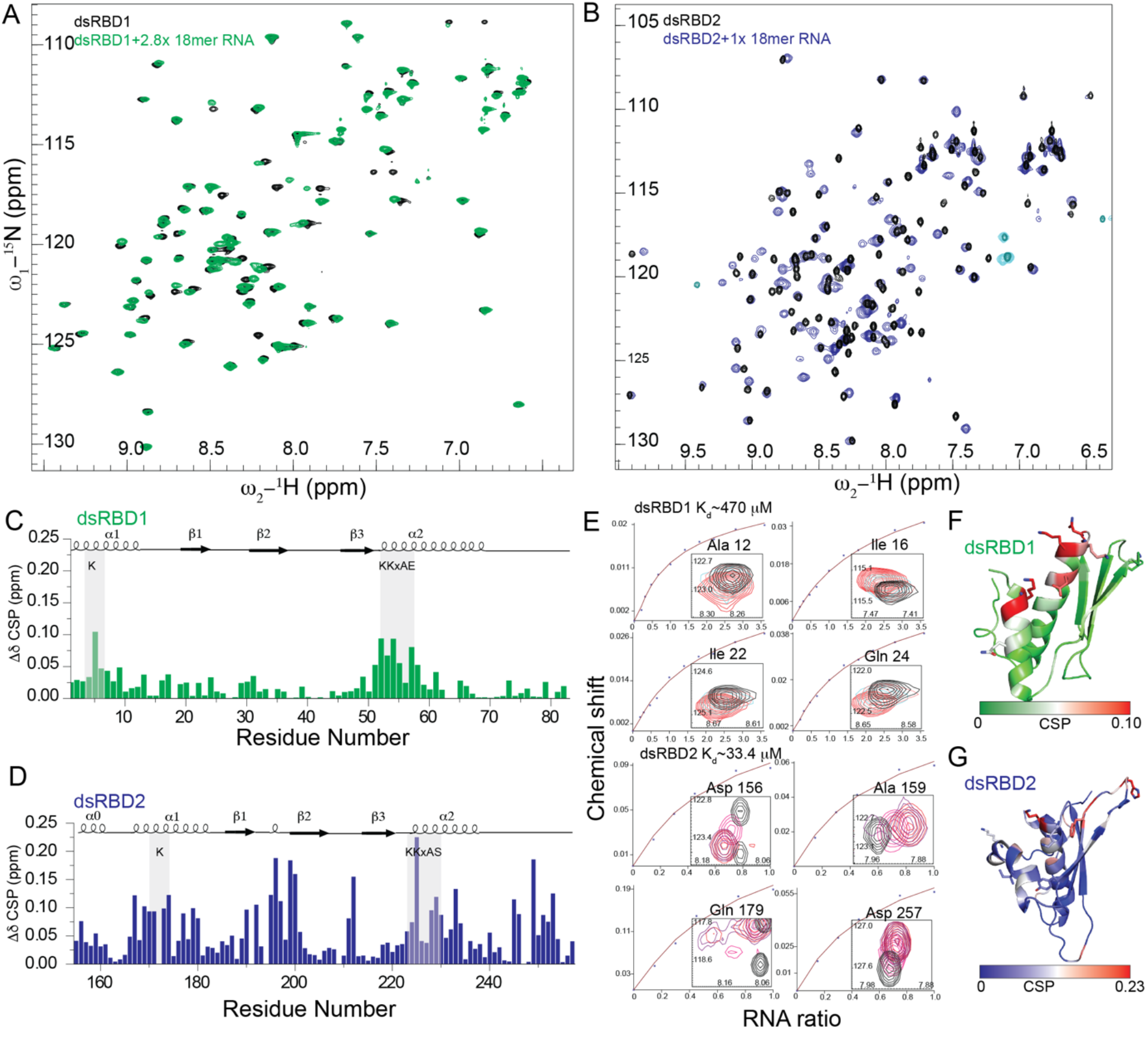
(A) & (B) Superposition of ^1^H, ^15^N HSQC NMR spectra of individual dsRBD1 and dsRBD2 domains free and bound to SL7^18mer^. Only the final titration point is shown here for clarity (see Figure S2 for all titration points). (C) & (D) Chemical shift perturbations upon titration with saturating concentrations of SL7^18mer^ in dsRBD1 and dsRBD2 respectively. (E) Fitted NMR titration curve along with the respective ^1^H,^15^N-HSQC peak shifts corresponding to the same residue. dsRBD1 and dsRBD2 showed an affinity of 470 µM and 33 µM respectively for SL7^18mer^. (F) & (G) Mapping of chemical shift perturbations on the NMR structure of dsRBD1 and dsRBD2, respectively.

Both dsRBDs showed clear binding to SL7^18mer^ (**Figure 4A, B**). RNA binding to dsRBD1 is in the fast exchange regime on the NMR chemical shift time scale whereas for dsRBD2 some residues show RNA binding in the intermediate exchange regime, (**Figure 4E**) where peaks disappear at a low molar ratio of dsRNA and reappear upon equimolar addition of dsRNA (eg. Gln179). This indicates that dsRBD2 has a higher affinity to RNA than dsRBD1. Fitting the NMR chemical shift perturbations for several residues showing fast exchange behavior vs RNA concentration yielded an average dissociation constant of ~470 µM and ~30 µM for dsRBD1 and dsRBD2, respectively, confirming that amongst the two domains, dsRBD1 binds RNA with high µM affinity and thus has binding weaker than dsRBD2. The affinity of dsRBD2 is in the lower µM range which is typical for single dsRBD domains. Given that dsRBD2 shows stronger chemical shift perturbations (**Figure 4C, D**) and has a higher affinity for double stranded RNA binding compared to dsRBD1 suggests that dsRBD2 could potentially drive the binding of RNA to MLE also in the full-length context. Plotting the chemical shift perturbation on the NMR structure of dsRBD1 and dsRBD2 demonstrates that the majority of residues involved in dsRBD-RNA interactions are limited to α_1_, α_2_, β_1_ and the loop connecting β_1_ and β_2_ sheets (**Figure 4 F, G**), in agreement with the identification of RNA binding residues typical for dsRBD domains.

### Interaction of dsRBD1,2 with SL7 roX2 RNA

To understand the binding of individual dsRBD domains in the context of dsRBD1,2 and to study if there is any cooperativity between the two domains for RNA binding, we performed filter binding assays and ^1^H,^15^N-HSQC NMR titrations of dsRNA into tandem dsRBD1,2 domain (**Figure S5**). As determined from filter binding experiments, the affinity of dsRBD1,2 for SL7 binding was a modest 2-fold higher than the dsRBD2 alone suggesting an additive effect between the two domains (**Figure S5D**). For deciphering the mode and specificity of dsRBD1,2 for roX2 binding, we performed ^1^H,^15^N-HSQC NMR titrations with 4 different dsRNAs; three derived from roX2 SL7 (cognate substrate of MLE; SL7^18mer^, SL7^23mer^ and SL7^14mer^Loop) and one derived from an unrelated dsRNA; UR^23mer^ (**Figure S4**). We varied the length of roX2 SL7 RNA to study the effect of RNA length on the binding affinity of dsRBD1,2-roX2 interaction (SL7^18mer^ and SL7^23mer^) and designed a minimal SL7 RNA consisting of the SL7 roX-box region fused to the apical stem-loop of SL7 (SL7^14mer^Loop) to study whether the SL7 apical loop has any effect on dsRBD1,2 interactions.

Titration of dsRBD1,2 with double stranded SL7^18mer^, SL7^23mer^ and UR^23^mer led to severe line broadening with increasing concentrations of RNA. At equimolar or slight excess of RNA concentrations, only peaks corresponding to linker residues remained visible (**Fig S5A-C**). As the exchange-broadened peaks of dsRBD1,2 do not reappear at excess concentrations of dsRNA, it is likely that the two domains slide over the dsRNA length as observed before in the case of TRBP and Loqs-PD (25,48). Thus, for these RNAs we cannot determine an affinity by NMR. Furthermore, as both, specific and non-specific RNA’s lead to similar line broadening effects in ^1^H,^15^N-HSQC spectra, we conclude that dsRBD1,2 does not show any specificity towards the three different RNAs.

As both domains in the tandem dsRBD1,2 showed line broadening upon binding to SL7^18mer^, SL7^23mer^ and UR^23mer^ likely due to the sliding motion of the protein along the RNA, we wondered whether the presence of an apical loop as found in the cognate SL7 substrate of MLE at one end of the dsRNA along with a slightly shorter RNA would alter the binding properties of dsRBD1,2 to RNA and circumvent line broadening in ^1^H, ^15^N HSQC NMR titration experiments. To test this, we performed titrations with SL7^14merLoop^ dsRNA obtained by fusing the SL7 roX-box with its apical loop. Here, a few peaks exhibit line broadening (e.g Lys 4, Gly 250) (**Figure 5A, S6**) upon addition of slight excess concentration of SL7^14merLoop^, suggesting only minor or no sliding motion of dsRBD1,2. The RNA binding of individual dsRBD domains within the tandem dsRBD1,2 construct is in the fast to intermediate exchange range suggesting that both domains now bind RNA with slightly higher affinity than the individual domains owing to cooperativity between the two domains in the dsRBD1,2 tandem construct. This is further confirmed by filter binding experiments (**Figure S5D**) wherein dsRBD1,2 binds SL7^14merLoop^ with 6.8 µM compared to the individual dsRBD2 domain which binds with an affinity of ~22 µM to SL7^14merLoop^ and dsRBD1 alone showed very weak binding to SL7^14merLoop^, in line with NMR titrations.

**Figure 5.**
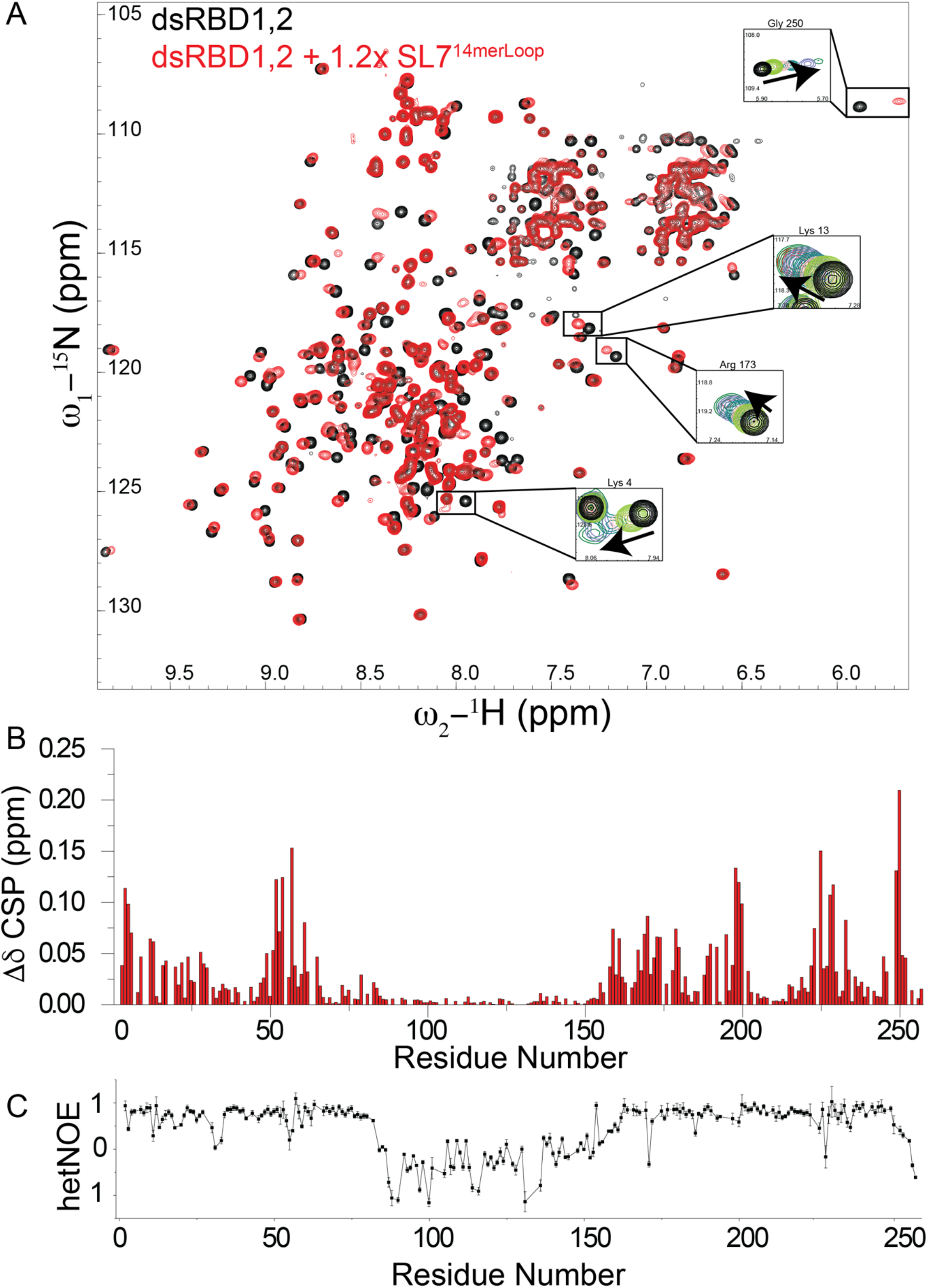
(A) Comparison of ^1^H,^15^N-HSQC NMR spectrum of dsRBD1,2 free (black) and bound to excess concentrations of SL7^14merLoop^ (red). Full titration points for some of the residues are shown in the insets and Figure S6. (B) Chemical shift perturbations in dsRBD1,2 upon titration with SL7^14merLoop.^(C) {^1^H}–^15^N heteronuclear NOE analysis of dsRBD1,2 bound to SL7^14merLoop^. Error bars are calculated from the spectral noise.

### Linker between dsRBD1,2 domains does not contribute to RNA binding

Several RNA binding proteins containing tandem RNA binding domains connected by flexible linkers show active participation of the linker residues for RNA recognition (49-52). In these proteins, the linker, which is flexible in the unbound state, adopts a folded conformation in the RNA bound state. As dsRBD1,2 is connected by a 95 amino acid long, flexible linker and the tandem construct shows higher affinity for SL7 binding than the individual domains (**Figure S5D**), we wondered whether the linker contributes to the binding of roX2 RNA *in vitro* and adopts a preferred conformation or secondary structure upon RNA binding. We did not observe any significant chemical shift perturbations upon RNA titration in tandem dsRBD1,2 (**Figure 5B**). As many residues in individual dsRBD1 and dsRBD2 showed line broadening upon RNA titration, we also compared the line broadening in the free and RNA bound form of dsRBD1,2. If linker residues would contribute to RNA binding, we would expect a significant drop in intensity of the corresponding resonances due to reduced overall tumbling upon RNA binding. However, the linker resonances show only a slight decrease in peak intensity compared to the two domains which show a large drop in the intensity. To further test if the linker adopts any structured conformation in the RNA bound state of dsRBD1,2, we recorded {^1^H}–^15^N heteronuclear NOE experiments (**Figure 5C**). As this region still showed complete flexibility in these experiments, with no chemical shift perturbations in the linker region upon dsRNA titration and its retained high flexibility we can conclude that the linker is not involved in RNA binding.

To test more rigorously and to identify regions of the linker, if any, which could contribute to RNA binding, we created a linker deletion mutant, where a large part (A85-I140) has been removed. We determined the RNA binding activity of this mutant to roX2 SL7 RNA and SL7^14merLoop^. Filter binding experiments showed that deletion of the linker region did not lead to any significant effect on the binding affinity of dsRBD1,2 to both RNAs (**Figure S5D**) further supporting our NMR observation that the linker does not contribute to RNA binding and is in fact dispensable.

### Mutations of dsRBD1 affect RNA binding in the context of full-length MLE

To validate the observed dsRBD1-RNA interactions, we mutated RNA binding residues (K4E, K53E, K54E and K53+54E) in dsRBD1, which exhibited the strongest chemical shift perturbations in our NMR titration experiments (**Figure 4**). To determine whether the mutants bind to dsRNA, we performed ^1^H,^15^N-HSQC NMR titration of ^15^N labelled dsRBD1 mutants with SL7^18mer^ dsRNA (**Figure S7A-E**).Spectra of K53E, K54E and K53E+K54E mutants in presence of excess concentrations of dsRNA were identical, concluding that these mutants do not bind dsRNA. Only K4E showed minor dsRNA binding at 1.3x excess of SL7^18mer^ but has strongly impaired dsRNA binding properties. We also confirmed dsRNA binding of dsRBD1 mutants by filter binding experiments where none of the mutants showed significant RNA binding compared to wild type dsRBD1 (**Figure S7F**). As dsRBD2 showed significantly stronger binding than dsRBD1 and other domains in MLE including the G-paτ_c_h could potentially bind RNA and obscure the contribution of dsRBD1 for RNA binding in the dsRBD1,2 and MLE full-length (MLEfl) context, we also evaluated the binding of dsRNA to dsRBD1 mutants in the context of dsRBD1,2 (**Figure S8A**) and MLEfl by filter binding assays (**Figure 6A**). In the dsRBD1,2 construct, mutations of the dsRBD1 domain resulted in ~2- to 4-fold reduction in RNA binding. The dsRBD2 mutant K225E showed a severely diminished but still detectable RNA binding. The additional mutation of K54E led to complete loss of RNA binding in the dsRBD1,2 K54+225E mutant. In the context of MLEfl, the RNA binding mutations impaired roX2 binding. Similar to the mutations in the dsRBD1,2 construct, K225E did show very little RNA binding while the K54+225E double mutant did not bind RNA at all, confirming the contribution of dsRBD1 to RNA binding *in vitro*.

**Figure 6.**
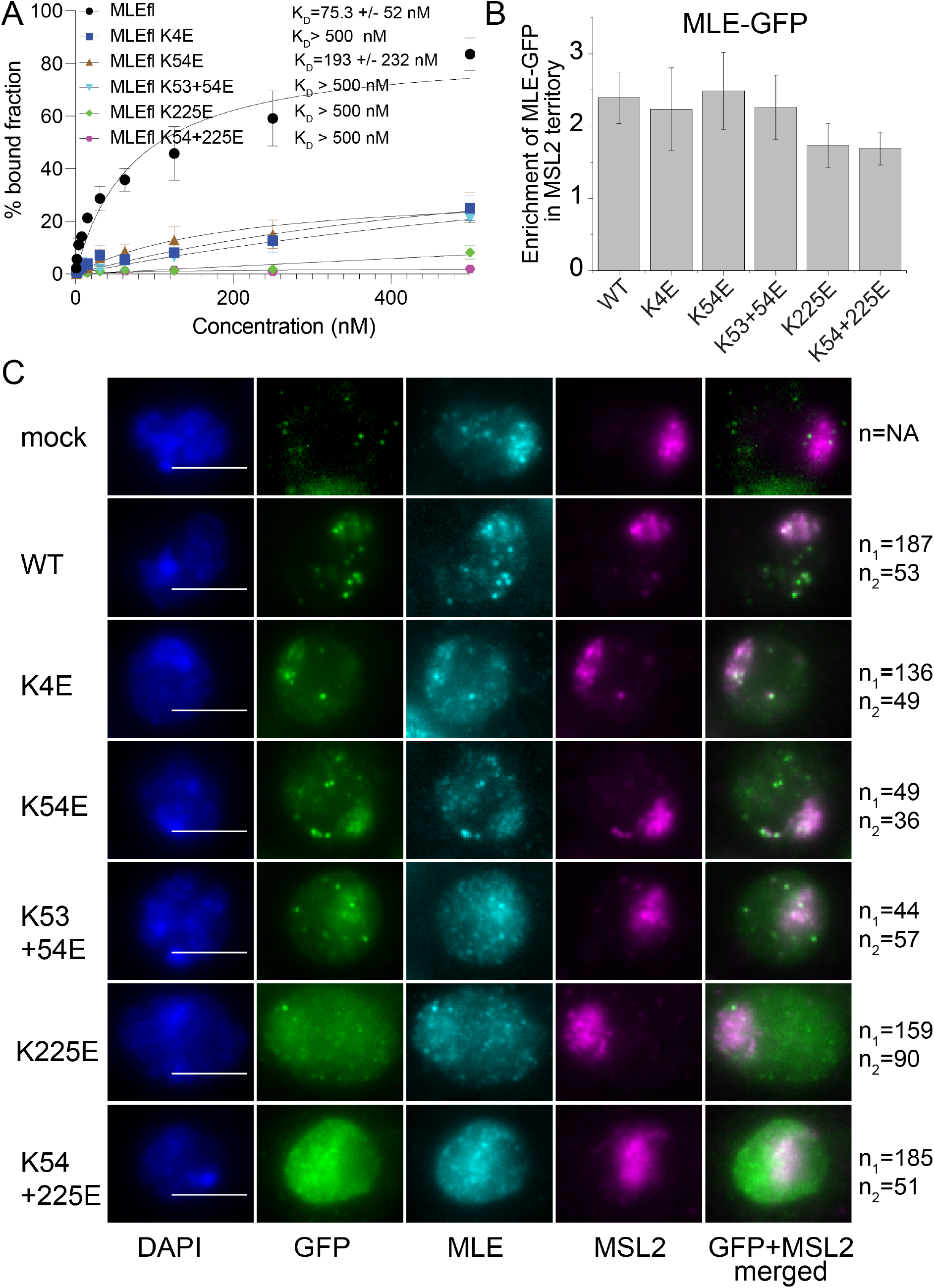
(A) Filter binding experiments with MLEfl-dsRBD mutants. Error bars represent standard deviation of two experiments. (B) Enrichment of GFP-MLE and its mutants on X chromosome territory is shown. Error bars represent standard deviations for two independent biological replicates. (C) Representative images of analyzed cells for quantification of enrichment of GFP-MLE in X chromosome territory is shown. DNA counterstaining with DAPI and immunostaining against GFP, MLE and MSL2 along with the merged image for GFP and MSL2 channel is shown. n_1_ and n_2_ are the number of cells analyzed in two independent biological replicates respectively. The white scale bar in the DAPI channel image represents 5 µm.

### Effect of dsRBD RNA binding mutants on *in vivo* roX2 binding and MLE localization

As we show that dsRBD1 binds RNA *in vitro*, we wished to assess whether this interaction was relevant to known *in vivo* functions. We analyzed two functions of MLE in cells: the binding of MLE to its major substrate, roX2 RNA, and the localization of the helicase to the male X chromosome, which is known to depend on roX RNA (18,53-55). To this end, we established stable S2 cell lines expressing GFP-tagged wild type MLE and MLE dsRBD1/dsRBD2 domain mutants in the background of endogenous MLE. To test the *in vivo* binding of dsRBD1,2 mutants to roX2, we performed native RNA immunoprecipitation experiments (RIP) where GFP-tagged MLE derivatives were immunoprecipitated from whole cell extracts using an anti-GFP antibody. The fraction of bound roX2 was quantitatively determined by RTPCR using primers targeting either the 5’ end (SL3) or the 3’ end (SL7) of roX2. Extracts from nontransfected S2 cells (mock) served as negative control excluding non-specific sticking of RNA to beads. The analysis was complicated by the fact that dsRBD mutants were stronger expressed than wild type MLE after the required expansion of the stable cell lines, precluding a direct comparison of mutant and wild type helicase. Previously, the dsRBD2 mutant K225E showed severely diminished roX2 binding in an analogous experiment (18). In the current experiment (involving newly generated stable lines) this mutation also shows reduced roX2 binding and thus serves as a reference (**Figure S8D**). The additional mutation of K54E did not lead to further decreased roX2 binding. Furthermore, MLE mutated in K54E alone (which was expressed to similar levels as K225E, (see **Figure S8C**), or in combination with K53E immunoprecipitated roX2 much better. We conclude that the main determinants of roX2 binding in cells are located in dsRBD2, in agreement with earlier work (17,18). (6)

A functional MSL complex localizes to the X chromosome territory and can be easily identified by MSL2 immunostaining (41). The colocalization of MLE with this territory depends on the interaction with roX. Previously it had been shown that deletion of dsRBD1 partially impairs MLE binding to the X chromosome. To determine if this function of dsRBD1 depended on the RNA binding activity of dsRBD1, we assessed X territory localization of the MLE-GFP derivatives in the above-mentioned cell lines. As before, the dsRBD2 domain mutant K225E served as a negative control since this mutant fails to localize faithfully to a coherent X chromosomal territory, but rather displayed diffuse nuclear staining (18). These experiments were done during the initial phase of stable cell line selection, when the expression levels of MLE derivatives were still similar to each other and to endogenous MLE (**Figure S8B**). The localization of MLE-GFP and its mutants was assessed using antibodies targeted against GFP, MLE and MSL2. Staining of endogenous MLE served as an internal control for assessing the total cellular MLE amount and MSL2 staining identified X chromosome territories in the male S2 cells (41).

Automated analysis of X chromosome localization relative to the staining of MLE-GFP in the remainder of the nucleus, revealed that wild type MLE-GFP was enriched 2.4-fold in the MSL2 territory (**Figure 6 B,C**). A comparable enrichment was observed for all dsRBD1 mutants. In contrast, the X chromosome enrichment of dsRBD2 mutant K225E was impaired. Therefore, our analysis suggests that at least at a quantitative level and in presence of endogenous MLE, RNA binding through dsRBD1 is not required for *in vivo* X chromosome localization of MLE-GFP.

## DISCUSSION

In this study, we identify the dsRBD1 domain as a *bona fide* dsRNA binding domain of MLE and report the high-resolution structure of dsRBD1 and dsRBD2 within a dsRBD1,2 tandem construct. The two dsRBD domains in absence of RNA tumble as independent modules as shown by absence of any interdomain NOEs, relaxation analysis and lack of chemical shift perturbations between the isolated domains and the tandem construct.

Our solution NMR structure of the dsRBD2 domain in dsRBD1,2 context superimposes well with the dsRBD2 structure present in the crystal structure of the MLE core region. Interestingly, in the crystal structure, the helix α0 in dsRBD2 makes multiple charged and van der Waals interactions with residues from helix α1 of dsRBD2, the loop connecting β hairpin 1 and 2 of the RecA2 domain and the loop connecting β4 and β5 loop of the auxiliary OB-like fold (18) (**Figure S2B**). Residues from β4-β5 loop of auxiliary OB like fold are directly involved in specifically recognizing the uracil base of poly-U RNA used in the study. Therefore, packing of α0 in dsRBD2 could potentially indirectly affect the interaction of MLE with roX2 RNA in the full-length context. However, in our NMR structure, helix α0 is completely flexible and does not form any specific interaction with α1. This could possibly be due to the absence of interactions from other domains which are present in the full-length context. Nevertheless, we did not observe chemical shift perturbations and thus any interaction of α0 helix with dsRNA in our NMR titration experiments, negating its role in direct protein-RNA interaction for the binding of MLE to double stranded roX2 RNA. A similar α0 helix has been observed in the dsRBD1 domain of the TRBP + siRNA complex structure where also no interaction of this helix with siRNA was detected (24).

Previous structural and biochemical studies had only identified the role of dsRBD2 and G-paτ_c_h for dsRNA binding (17-20). The role of dsRBD1 in dsRNA binding was not clear due to no or just minor binding observed in electro mobility shift assays (EMSA)(9,19). However, as protein-nucleic acid complexes are not at binding equilibrium during EMSA experiments, probing protein-RNA interactions with fast dissociation constants could pose a significant challenge and could lead to false negatives (56). In NMR titration experiments with individual dsRBD domains, on the other hand, we could clearly observe RNA binding to both dsRBD domains. Interestingly, both dsRBDs exhibit a non-canonical KKxAK motif, where an additional residue, a glutamate (at position 57) in dsRBD1 and a serine (at position 228) in dsRBD2 (KKxAE and KKxAS motif in dsRBD1 and dsRBD2 respectively) replaces the canonical last lysine. An additional lysine follows immediately in the sequence after KKxAE/S in both domains, however, it might not be able to make contacts with RNA as the additional E/S in the sequence of both domains spaces it further apart from the RNA binding site. A similar situation has been observed in RNase III and DHX9 dsRBDs, where in absence of the third lysine in the KKxAK motif is complemented by a lysine from helix α1 at position 3 (21,57). In this case, recognition of the RNA major groove is mediated by a bipartite motif where the first two lysines come from the KKxAK motif and the third lysine comes from helix α1. In the MLE dsRBD1,2 structure presented here we also observe a third lysine optimally oriented for dsRNA recognition (located at position 4 in dsRBD1 and the corresponding position 171 in dsRBD2) which could potentially rescue the incomplete KKxAK motif. Mutation of two lysines from the KKxAE motif led to complete loss of RNA binding by dsRBD1. Although, the K4E mutant retained minor RNA binding activity as detected by NMR titrations for the isolated domain, it bound even weaker compared to the K54E mutant in the full length and tandem dsRBD1,2 context signifying its importance for functional complementation of the non-canonical KKxAK motif in the MLE dsRBD1,2.

^1^H, ^15^N HSQC NMR titration of dsRBD1,2 with 18 and 23 bp long double stranded RNA’s derived from stem loop 7 (SL7) of roX2 led to severe signal loss. This strongly suggests sliding of dsRBD1,2 on these RNAs as changing chemical environment in the dynamically exchanging protein-RNA complex leads to line broadening. Similar sliding motions have previously been observed for several other dsRBDs and have been shown to be a prerequisite for the biological activity of some (46,48,58). As roX2 consists of eight stem loops, it will not be surprising that MLE initially utilizes the sliding mechanism by engaging its dsRBD1,2 domains to find an optimal site for initiating helicase activity to unwind the SL7 roX-box region for binding of MSL2. Sliding of dsRBD1,2 on SL7^18mer^ and SL7^23mer^ RNA’s also suggested a possibility of dsRBD1,2 recognizing non-specific dsRNA sequences as recognition of specific sequences by dsRBDs will lead to tight binding to one specific RNA region and thus stabilization of a single conformation of a dsRBD1,2+RNA complex. Indeed, titration with an unrelated UR^23mer^ double-stranded RNA showed a similar mode of binding to MLE dsRBD1,2 along with severe line broadening suggesting again sliding of dsRBD1,2. Thus, we propose that dsRBD1,2 bind double-stranded roX2 RNA in a structure-specific but not sequence-specific manner. Given that MLE specifically recognizes stem structures containing U-rich roX-box sequences (9-11), the sequence specificity to U-rich single stranded sequences must come from the helicase core of MLE as has been shown before (18).

^1^H, ^15^N HSQC NMR titration experiments with a slightly shorter version of SL7 roX-box region capped at one end with the apical loop of SL7 (SL7^14merLoop^) to prevent sliding of dsRBD1,2 on the RNA showed minimal line broadening. As both domains showed simultaneous binding to SL7^14merLoop^ RNA as confirmed by chemical shift perturbations in both domains upon SL7^14merLoop^ titration and filter binding experiments, we can conclude that both domains in the tandem construct bind to this RNA. Binding of two tandem dsRBD domains to short sequences similar to SL7^14merLoop^ have been shown before for nuclear factor 90 (NF90) dsRBDs wherein the two domains bind on either side of the short 18mer duplex RNA and show a binding stoichiometry of 1:2 RNA to protein (59). Therefore, dsRBD1,2 could indeed be accommodated on either side of the double stranded SL7^14merLoop^ RNA and as there is no free RNA surface remaining for further interactions with either dsRBDs, this could prevent sliding of the dsRBD1,2 on the SL7^14merLoop^ as observed in our NMR experiments. In all the NMR titrations with SL7^18mer^, SL7^23mer^, UR^23mer^ and SL7^14merLoop^, we only observed minor line broadening and no chemical shift perturbations in the linker residues suggesting that the linker region between dsRBD1 and dsRBD2 in the tandem construct does not contribute to RNA binding and remains flexible; a fact further confirmed by NMR {1H}–^15^N heteronuclear NOE experiments of the dsRBD1,2+ SL7^14merLoop^ complex (**Figure 5C**). Importantly, the linker is completely dispensable for RNA binding as demonstrated by filter binding assays for SL7, where a linker deletion construct binds with affinities similar to wild type dsRBD1,2. Also, binding of shorter SL7^14merLoop^ is not affected between wild type and linker deletion.

In this study, we show that dsRBD1 is indeed involved in RNA binding *in vitro*. We observe that mutations of the RNA-binding interface of dsRBD1 do not affect MLE binding to the X-chromosome suggesting that the RNA binding property of dsRBD1 is dispensable for MLE localization, at least at the resolution afforded by immunofluorescence microscopy. Furthermore, since we analyzed MLE-GFP localization in steady state of stable cell lines and in presence of endogenous MLE, we cannot exclude an effect of dsRBD1 on the kinetics and efficiency of MSL complex assembly at the early stages of the establishment of dosage compensation in *Drosophila* embryogenesis, when MLE is found mainly associated with maternal roX1. Such a role in fine-tuning of MSL complex assembly may still involve RNA binding. Both dsRBDs are connected by the long, flexible linker and upon tethering of MLE through dsRBD2, dsRBD1 can ‘search’ for additional RNA targets by a kind of fly casting mechanism.

Other roles of dsRBD1 may also be envisaged. MLE has additional functions beyond dosage compensation, for example in RNA editing (16) and processing of siRNA precursors (15). In such a context, dsRBD1 may synergize with dsRBD2 for specialized functions. Furthermore, dsRBDs have recently been shown to be involved in protein-protein interactions, including homodimerization (60-62). Further studies will help to unravel the specific role of dsRBD1 in MSL localization and to identify cofactors, e.g. Unr (11), potentially required in association with dsRBD1 for MSL assembly.

## DATA AVAILABILITY

Atomic coordinates for the dsRBD1,2 NMR structure have been deposited with the PDB under accession number 6I3R. Chemical shifts and NMR data for structure determination of dsRBD1,2 have been deposited to the BMRB under accession number 34326.

## SUPPLEMENTARY DATA

Supplementary Data is available

